# Ykt6 SNARE protein drives GluA1 insertion at synaptic spines during LTP

**DOI:** 10.1101/2025.02.10.632800

**Authors:** Momoko Takahashi, Gabriela Caraveo

## Abstract

Long-Term Potentiation (LTP), a crucial form of synaptic plasticity essential for memory and learning, depends on protein synthesis and the upregulation of GluA1 at postsynaptic terminals. While extensive research has focused on the role of endosomal trafficking in GluA1 regulation, the contribution of endoplasmic reticulum (ER) trafficking pathways remains largely unexplored. A key opportunity to investigate this emerged from Ykt6, an evolutionarily conserved SNARE protein and a master regulator of vesicular fusion along ER-trafficking pathways. Here, we demonstrate that Ykt6 is highly expressed in the mammalian hippocampus, where it localizes to synaptic spines and regulates GluA1 surface expression in an LTP-dependent manner. Furthermore, we found that Ykt6 modulates synaptic vesicle pool dynamics as well as the amplitude and frequency of miniature excitatory postsynaptic currents. Ykt6 loss of function has been linked to α-synuclein pathology, a hallmark of Lewy Body Dementias (LBDs), where α-synuclein misfolding in the hippocampus disrupts LTP. Taken together, our findings establish Ykt6 as a critical SNARE protein in hippocampal function during LTP, with significant implications for neurodegenerative disorders such as LBDs.

## Introduction

Through an unbiased phosphoproteomic approach, our laboratory previously identified Ykt6 as a substrate of the Ca^2+^-dependent serine/threonine phosphatase, calcineurin (CaN) (1). Ykt6, an essential soluble N-ethylmaleimide-sensitive factor attachment protein receptor (SNARE), regulates vesicular fusion along the secretory pathway, namely the transport between the endoplasmic reticulum (ER) and Golgi apparatus, within the Golgi, from the Golgi to the plasma membrane (2, 3), and in autophagy-related vesicular fusion pathways (1, 4–10). Unlike most SNARE proteins, Ykt6 lacks a transmembrane domain and relies on reversible lipidation for recruitment to membranes to facilitate vesicular fusion. We and others have established that in the inactive state, Ykt6 is localized in the cytosol in a close conformation, whereby the regulatory longin domain is closely associated with the SNARE domain (1, 3, 10–12). In the active state, phosphorylation at the SNARE domain builds up the electrostatic potential which causes an intra-conformational change separating the longin and the SNARE domain enabling C-terminus lipid modifications (1, 11, 13–17). Reversible lipidation allows Ykt6 to be anchored to the ER, Golgi and plasma membranes (1, 13, 14, 16, 18, 19). Our laboratory also discovered that the subsequent dephosphorylation of the SNARE domain by CaN enhances SNARE-SNARE protein interactions, thereby facilitating vesicular fusion in both the secretory and autophagic pathways (1).

At excitatory synapses, α-amino-3-hydroxy-5-methyl-4-isoxazolepropionic acid receptors (AMPARs) modulate synaptic strength to facilitate information processing and storage (20–23). AMPARs are rapidly inserted into synapses during long-term potentiation (LTP), a form of synaptic strengthening (24). While the roles of exocytosis and endocytosis in activity-dependent AMPAR transport at postsynaptic terminals are well-established (25–29), little is known about the contribution of activity-dependent roles of the secretory pathway. GluA1, an AMPAR subunit, is a good candidate for studying secretory transport in relation to synaptic plasticity in hippocampal neurons. First, GluA1 as the rest of AMPAR subunits, are synthesized in the ER where they assemble to form the channel (30–32). Second, GluA1A2 heteromers are the predominant AMPA receptor type that mediates synaptic transmission in the hippocampus (33–35). Third, while GluA2 localizes around the plasma membrane and remains relatively static, GluA1 relies more on local protein synthesis and activity-dependent trafficking for constant transport to the synaptic space (36–38). Lastly, synaptic insertion of GluA1-containing AMPARs is necessary for LTP (39–46).

LTP triggers Ca^2+^ influx through both N-methyl-D-aspartic acid receptors (NMDARs) and AMPARs, and our previous findings in yeast and HeLa cells demonstrated that Ykt6 activity is regulated by CaN (1), whose activity is dependent on Ca^2+^. Moreover, we and others have implicated Ykt6 loss of function in α-synucleinopathies, neurodegenerative diseases characterized by deficits in LTP (1, 47–49). Therefore, we asked whether Ykt6, the master regulator of the secretory pathway, participates in delivering GluA1 subunits at synaptic terminals during LTP. Here we show that Ykt6 is highly expressed in the hippocampus in the mammalian brain. Using primary pyramidal hippocampal neurons, we demonstrate that Ykt6 relocates to synaptic spines in response to LTP and promotes surface expression of GluA1. Moreover, we show that Ykt6 regulates the number of synaptic vesicular pools, the amplitude and frequency of miniature excitatory postsynaptic currents. Taken together, our findings highlight a critical role for Ykt6 in LTP with implications to α-synucleinopathies.

## Results

### Ykt6 is highly expressed in the mammalian hippocampus

We first began by examining Ykt6 expression in the mammalian brain using the human and mouse brain atlas (50, 51). In the human brain, data from six subjects who were otherwise healthy at the time of death was available. We examined Ykt6 expression using two different mRNA probes targeting Ykt6 in different brain regions, including the hippocampus. As a reference control, we analysed the expression of the neuronal specific microtubule associated protein 2 (MAP2), a highly expressed neuronal protein. Relative to MAP2, the highest expression of Ykt6 was in the globus pallidus followed by the hippocampus (Figure 1A, hippocampal formation). In the rodent brain, while the mRNA probe for Ykt6 was ubiquitously expressed throughout the brain, its expression was highest in the hippocampus and cerebellum, followed by the frontal cortex (Figure 1B). Together, these data show that under physiological conditions Ykt6 is highly expressed in the mammalian hippocampus (52–64).

**Figure 1.**
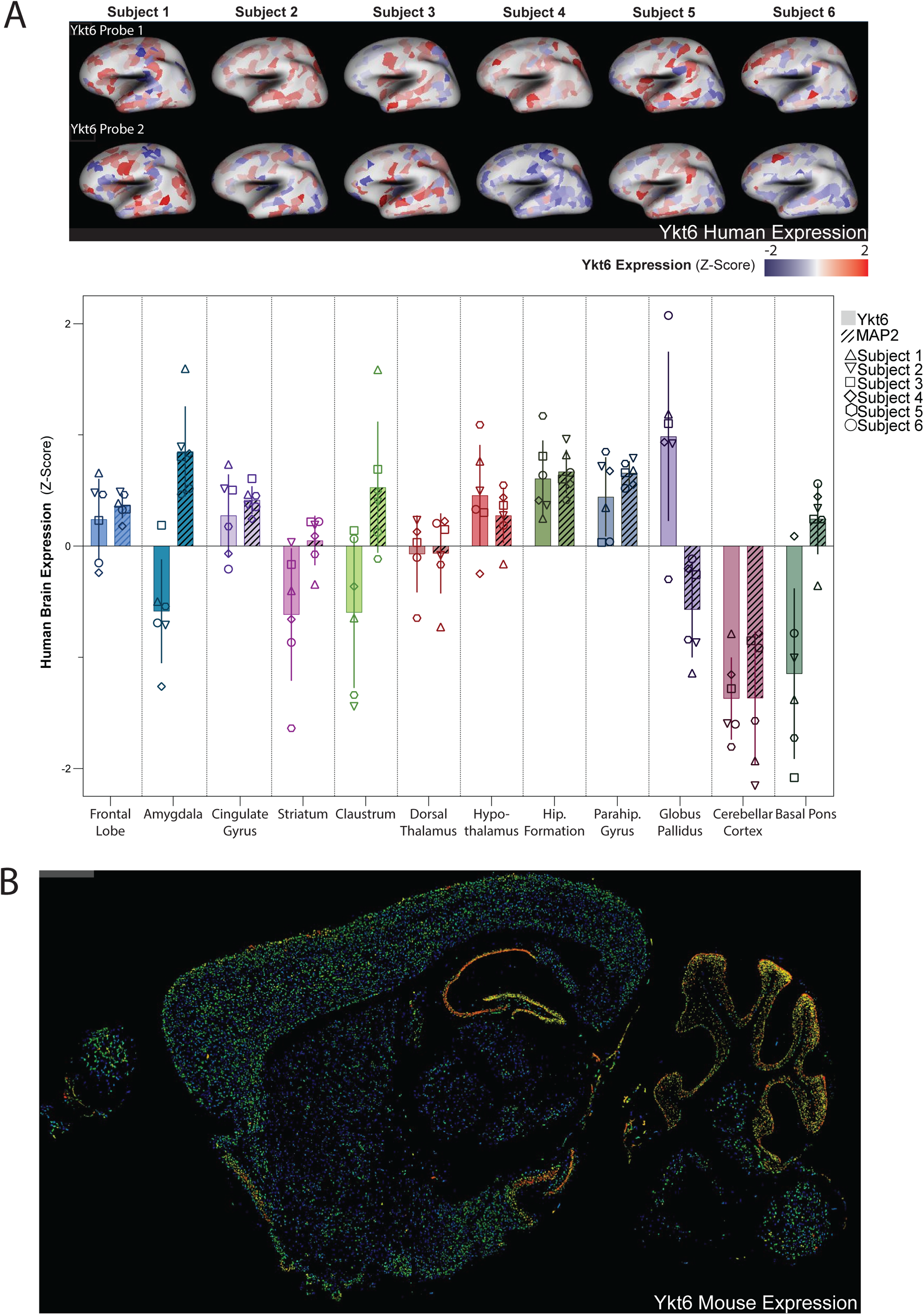
Ykt6 is highly expressed in the hippocampus in the mammalian brain. **A)** Top, Sagittal view of the Ykt6 expression by in-situ hybridization assay and microarray with two Ykt6 probes in healthy adult human brains. Allen Human Brain Atlas, https://human.brain-map.org/microarray/search/show?search_type=user_selections& user_selection_mode=2. Bottom panel, quantification of Ykt6 and Microtubule-associated protein 2 (MAP2) from the indicated brain regions (154). N=6 humans; MAP2 serves as a metric for high expression. Error bars represent standard deviation. **B)** Ykt6 expression by fluorescence in-situ hybridization in a sagittal section from a mouse brain. Allen Mouse Brain Atlas, https://mouse.brain-map.org/experiment/show/71380453.

### At rest, Ykt6 is mainly cytosolic, with some colocalization with the Golgi apparatus and ER at somatic and dendritic terminals in hippocampal neurons

We next examined the intracellular localization of Ykt6 in rat primary hippocampal neurons in culture using immunofluorescence. In resting conditions, Ykt6 was primarily cytosolic in both the somatic and dendritic regions (Figure 2A-D,I-L). Consistent with its role in the secretory pathway (65), Ykt6 also localized to the Golgi apparatus and to the ER, as shown by the colocalization with the Golgi marker GM130 and the ER-resident protein disulfide isomerase (PDI) (Figure 2A-D,I-L). Ykt6 localization is specific, as reducing endogenous Ykt6 expression with a lentivirus expressing an inducible ShRNA targeting Ykt6 to knock down its expression (Sh Ykt6) significantly diminished Ykt6 detection compared to neurons expressing a scramble ShRNA sequence as a control (Sh Ctrl) (Figure 2E-H,M-P). Together, these data indicate that under resting conditions, Ykt6 is primarily cytosolic but also associates with the ER and Golgi apparatus at somatic and dendritic locations.

**Figure 2.**
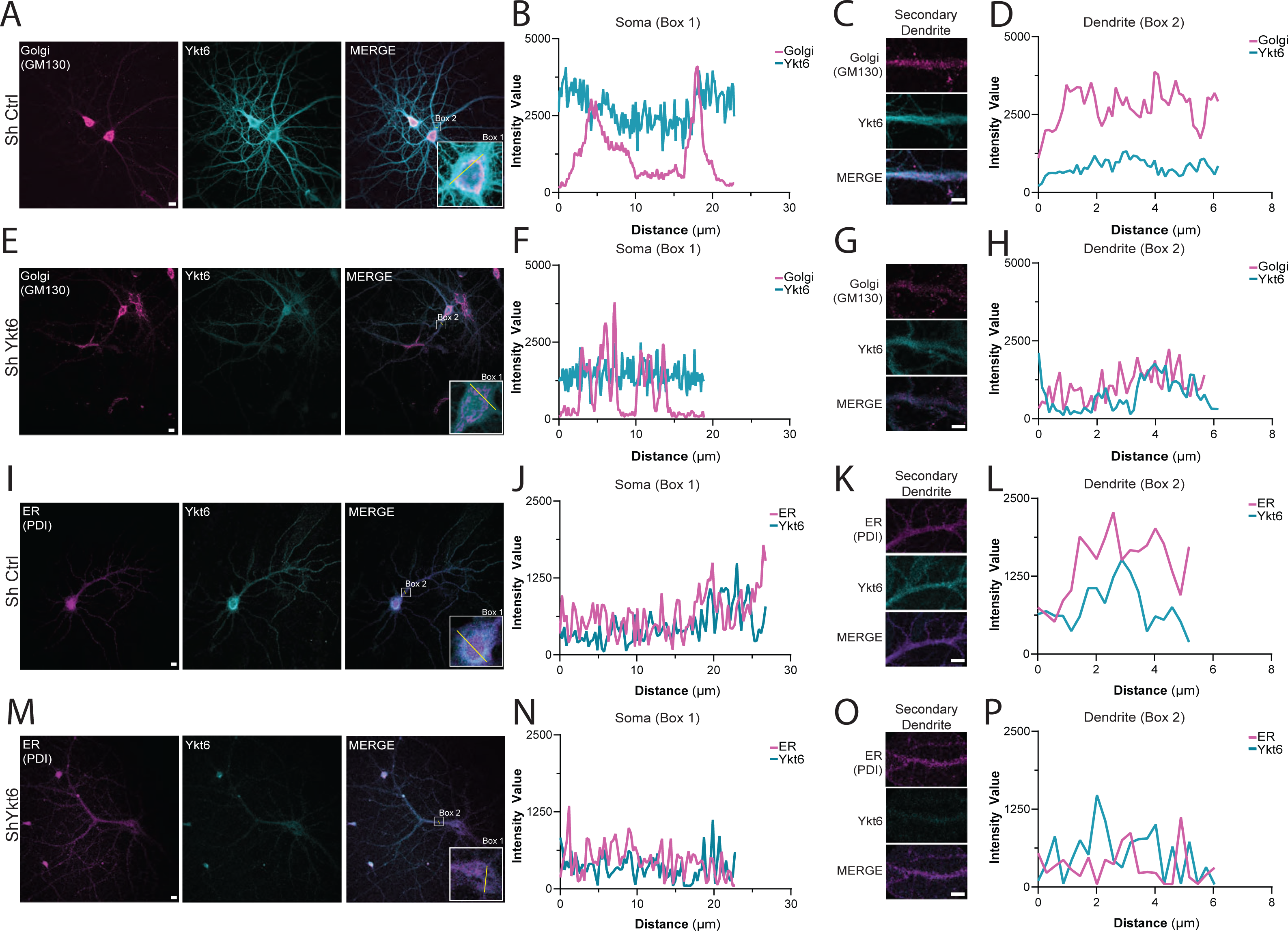
Ykt6 is present in the cytosol, Golgi Apparatus, and endoplasmic reticulum at somatic and dendritic locations in hippocampal pyramidal neurons. A-H) Primary hippocampal pyramidal neurons were transduced with either ShRNA against Ykt6 (Sh Ykt6) or ShRNA scrambled sequence as Control (Sh Ctrl) at DIV5, induced for Ykt6 knockdown with doxycycline starting at DIV8 and immunostained at DIV21 with GM130 (Golgi Apparatus) and Ykt6. **A)** Representative immunofluorescence images of neurons transduced with Sh Ctrl of Golgi apparatus (magenta) and Ykt6 (cyan); scale bar, 10µm. **B)** Fluorescence intensity linescans of Golgi apparatus and Ykt6 from the soma (Box 1) in (A). **C)** Representative immunofluorescence image of secondary dendrites from (A). Golgi apparatus shown in magenta, Ykt6 in cyan; scale bar, 10µm. **D)** Fluorescence intensity linescans of the secondary dendrites from Box 2 in (A). **E)** Representative immunofluorescence image of Golgi apparatus (magenta) and Ykt6 (cyan) from Sh Ykt6-transduced primary hippocampal pyramidal neurons; scale bar, 10µm. **F)** Fluorescence intensity linescans of Golgi apparatus and Ykt6 from the soma (Box 1) in (E). **G)** Representative immunofluorescence image of secondary dendrites from (E). Golgi apparatus shown in magenta, Ykt6 in cyan; scale bar, 10µm. **H)** Fluorescence intensity linescans of the secondary dendrites from Box 2 in (E). **I-P)** Primary hippocampal pyramidal neurons were transduced with either Sh Ykt6 or Sh Ctrl at DIV5, induced for Ykt6 knockdown with doxycycline starting at DIV8 and immunostained at DIV21 with protein disulfide isomerase (PDI) (ER) and Ykt6. **I)** Representative immunofluorescence images of neurons transduced with Sh Ctrl of ER (magenta) and Ykt6 (cyan); scale bar, 10µm. **J)** Linescan of fluorescence intensities of ER and Ykt6 from the soma (Box 1) in (I). **K)** Representative immunofluorescence image of secondary dendrites from (I). ER shown in magenta, Ykt6 in cyan; scale bar, 10µm. **L)** Fluorescence intensity linescans of the secondary dendrites from Box 2 in (I). **M)** Representative immunofluorescence images of ER (magenta) and Ykt6 (cyan) from Sh Ykt6-transduced primary hippocampal pyramidal neurons. **N)** Linescan of fluorescence intensities of ER and Ykt6 from the soma (Box 1) in (M). **O)** Representative immunofluorescence images of secondary dendrites from (M). ER shown in magenta, Ykt6 in cyan ; scale bar, 10µm. **P)** Fluorescence intensity linescans of the secondary dendrites from Box 2 in (M).

### Ykt6 mobilizes to synaptic spines in a cLTP-dependent manner

Next, we investigated whether Ykt6 is present at postsynaptic spines in hippocampal neurons. To address this, we analysed its localization using immunofluorescence relative to postsynaptic density protein 95 (PSD-95), a standard postsynaptic marker, and performed subcellular fractionation for biochemical assessment. At resting conditions, Ykt6 was present at postsynaptic spines as demonstrated by immunofluorescence through colocalization with the postsynaptic marker PSD95 (Figure 3A,B), and biochemically by its increased expression in the synaptosomal fraction (Figure 3C,D). We then asked whether the induction of chemical long-term potentiation (cLTP) with glycine, a widely used method to enhance synaptic plasticity at postsynaptic spines (66–71), affects Ykt6 localization. As reported by many others (72), cLTP treatment increased GluA1 expression at postsynaptic spines (Figure 3C,E). Interestingly, the induction of cLTP also increased Ykt6 expression at postsynaptic spines, as assessed by immunofluorescence, which showed colocalization with the postsynaptic marker PSD-95 (Figure 3A,B). This finding was further supported biochemically by its increased expression in the synaptosomal fraction (Figure 3C,D). Together, these data indicate that while Ykt6 is present at postsynaptic spines under resting conditions, it is further mobilized to postsynaptic spines upon cLTP induction.

**Figure 3.**
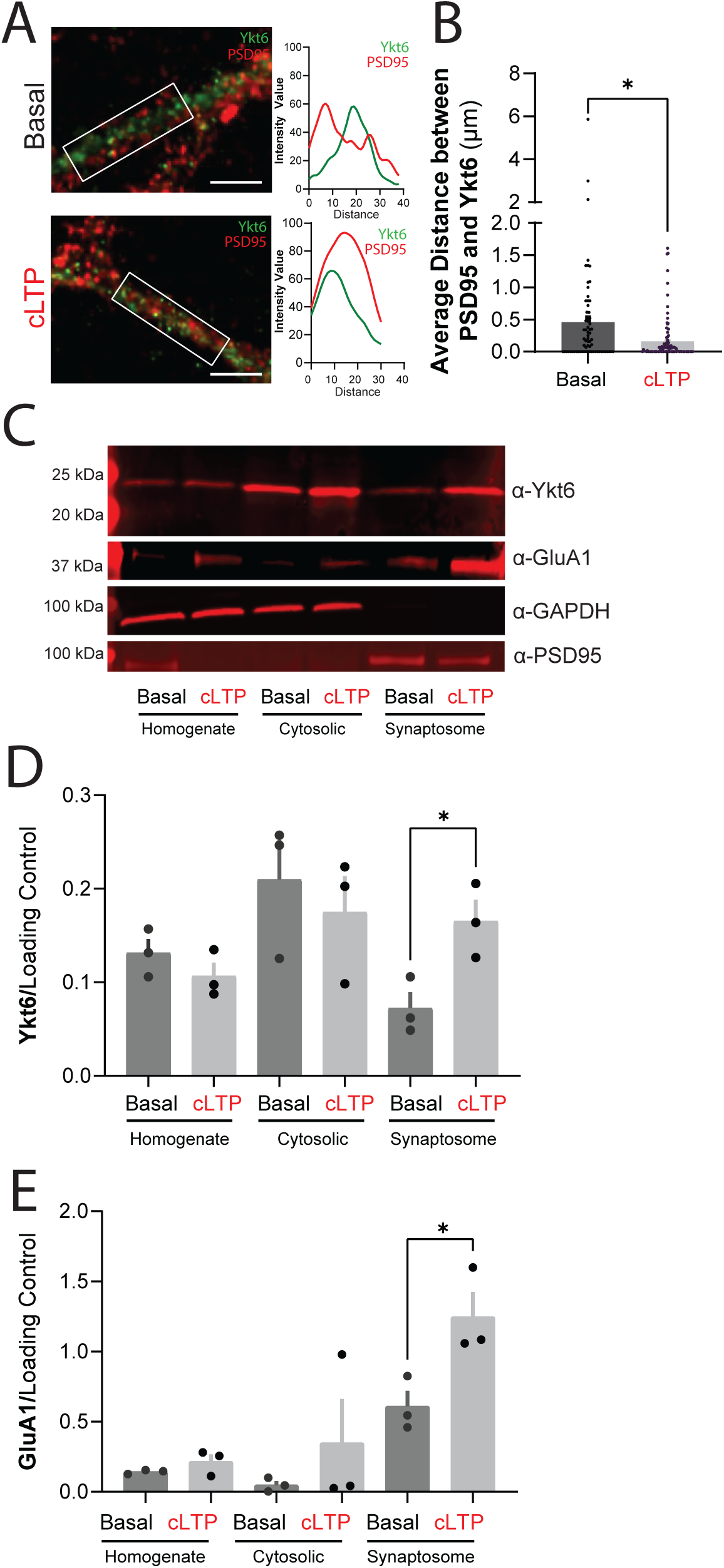
Ykt6 mobilizes to the postsynaptic spines in a cLTP-dependent manner. **A)** Left: representative images of secondary dendrites from primary pyramidal hippocampal neurons exposed to extracellular solution (ECS) as basal condition (top) or glycine for chemical long-term potentiation (cLTP, bottom) at DIV21 and immunostained with the postsynaptic marker, postsynaptic density 95 (PSD95) and Ykt6. Ykt6 in green, PSD95 in red. Scale bar, 5µm. Right: fluorescence intensity linescans from the region of interest (white box, left). (**B)** Quantification of average distance between PSD95 and Ykt6 from (A). Each data point represents aggregate average of all calculated shortest distances of each PSD95 puncta present in a selected ROI from a secondary dendrite to the closest Ykt6 puncta. N=2, 30-50 cells per biological replicate. Unpaired T-test. * p ≤ 0.05. **C-E)** Adult rat brain tissue was exposed to cLTP or ECS, fractionated to cytosolic and synaptic fractions and immunoprobed for Ykt6, GluA1, GAPDH and PSD95; GAPDH serves as a loading control for the homogenate and the cytosolic fractions, and PSD95 serves as a loading control for the synaptosomal fraction. Representative western blot (C) with quantification over respective loading controls of Ykt6 (D) or GluA1 (E). N=3. Unpaired T-test. * p ≤ 0.05.

### Ykt6 regulates GluA1 expression at synaptic spines in a cLTP-dependent manner

Modulation of the synaptic distribution of AMPARs underlies synaptic plasticity (20–23). In addition, AMPAR subunit assembly occurs within the ER (31, 32, 73, 74). Therefore, ER transport of AMPA receptors may modulate the synaptic pool available for membrane insertion, thereby impacting synaptic plasticity. Among the four AMPARs subunits (GluA1-4), GluA1 is a good candidate for studying secretory transport in relation to synaptic plasticity in hippocampal neurons. In the hippocampus, synaptic transmission is primarily mediated by GluA1A2 heteromers (33, 34). While GluA2 remains relatively stable near the plasma membrane, GluA1 depends on continuous intracellular trafficking to reach the synaptic space. Moreover, the synaptic insertion of GluA1-containing AMPARs is necessary for LTP (33, 39–46, 71, 75). To test if Ykt6 affects the distribution of endogenous AMPARs, we performed immunocytochemistry for surface and internal GluA1 on pyramidal neurons from the primary rat hippocampal cultures. The surface-to-internal ratio indicates that the regulation primarily occurs at the plasma membrane level rather than reflecting a global change in GluA1 expression. Primary hippocampal rat neurons were co-transduced with lentiviruses expressing an inducible ShRNA targeting Ykt6 to knock down its expression (hereafter referred to as Ykt6 knock down), or a scramble ShRNA sequence as a control (hereafter referred to as control). Additionally, an N-terminally green fluorescent protein (GFP) tagged human Ykt6 (GFP-Ykt6 WT), resistant to the Ykt6 ShRNA, was co-expressed to rescue Ykt6 expression to endogenous levels or GFP alone was used as control. Western blot analysis confirmed that Ykt6 knockdown reduces endogenous Ykt6 expression by approximately 70% and that the GFP-Ykt6 WT resistant construct successfully restores Ykt6 expression to endogenous levels (Figure 4A,B).

**Figure 4.**
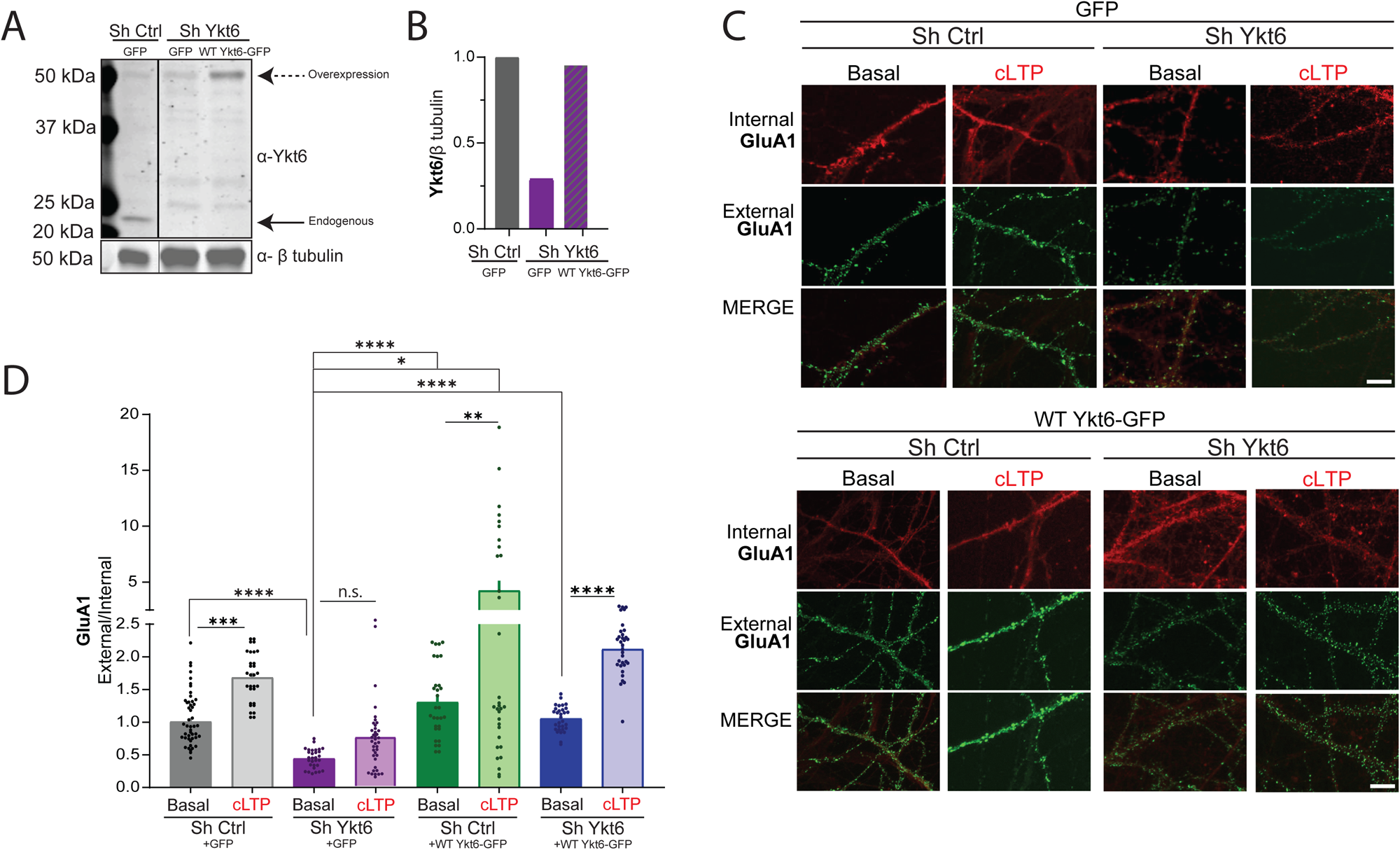
Ykt6 regulates GluA1 surface expression at synaptic spines in a cLTP-dependent manner. **A)** Representative western blot for Ykt6 expression from rat primary hippocampal neurons at DIV21, co-transduced with 3 different conditions: 1) Sh Ctrl + GFP, 2) Sh Ykt6 + GFP and 3) Sh Ykt6 + GFP-WT-Ykt6; solid arrow, endogenous Ykt6; dashed arrow, exogenous Ykt6. β-tubulin serves as a loading control. **B)** Quantification of Ykt6 endogenous expression over loading control from (A). **(C-D)** Rat primary hippocampal neurons co-transduced with 4 different conditions: 1) Sh Ctrl + GFP, 2) Sh Ctrl + GFP-WT-Ykt6, 3) Sh Ykt6 + GFP and 4) Sh Ykt6 + GFP-WT-Ykt6 were exposed to extracellular solution (ECS) or glycine for chemical long-term potentiation (cLTP) at DIV21 and immunolabelled for external and internal GluA1. **(C)** Representative immunofluorescence images of the secondary dendrites. Internal GluA1 in red, external GluA1 in green. Scale bar = 10µm. **(D)** Quantitation of ratios of external to internal GluA1 levels from (C), normalized to the control (basal condition, co-transduced with Sh Ctrl and GFP construct). One-way ANOVA with Welch’s correction and Dunnett’s multiple comparisons test. * p ≤ 0.05, ** p ≤ 0.01, *** p ≤ 0.001, **** p ≤ 0.0001. N=3, 8-10 cells per replicate.

Under resting conditions, Ykt6 knockdown slightly reduced surface expression of GluA1 compared to the control (Figure 4C,D). Overexpression of GFP-Ykt6 WT however, did not further increase surface GluA1 levels under basal conditions compared to control condition (Figure 4C,D), suggesting that GluA1 regulation is tightly controlled at rest. The effect on GluA1 surface levels under basal conditions was specific to Ykt6, as co-expression of a Ykt6 knockdown-resistant construct restored basal surface GluA1 levels (Figure 4C,D). As previously reported (72), cLTP treatment led to a two-fold increase in GluA1 surface expression under control conditions (Figure 4C,D, grey bars). Conversely, Ykt6 knockdown prevented the cLTP-induced increase in GluA1 surface expression (Figure 4C,D purple bars). Importantly, this effect was specific to Ykt6, as restoring endogenous Ykt6 expression with a knockdown-resistant construct reinstated the two-fold increase in surface GluA1 levels following cLTP treatment, similar to control conditions (Figure 4C,D blue bars). While overexpression of GFP-Ykt6 WT further increased surface GluA1 levels compared to the control under cLTP conditions, this effect was not statistically significant (Figure 4C,D green bars). Collectively, these findings indicate that Ykt6 regulates both basal and cLTP-dependent surface GluA1 levels.

### Ykt6 modulates both pre- and post-synaptic terminals of glutamatergic neurotransmission

Given the evidence that Ykt6 modulates synaptic connections and their downstream effects, we conducted whole-cell patch-clamp electrophysiology to measure miniature excitatory post-synaptic currents (mEPSCs) in pyramidal hippocampal neurons, providing further insight into synaptic dynamics. Rat primary pyramidal hippocampal neurons transduced with Sh Ctrl or Sh Ykt6 were patched and recorded in the presence of NMDA receptor blocker D-2-amino-5-phosphonopentanoic acid (D-APV), voltage-gated sodium channel blocker tetrodotoxin (TTX), and Gamma-Aminobutyric acid (GABA) subunit A (GABA_A_) receptor antagonist picrotoxin (PTX) to isolate AMPA-mediated currents. Ykt6 knockdown reduced the amplitude of mEPSCs compared to the control without significantly changing the distribution of the mEPSC amplitudes (Ykt6 Knockdown, 8.728 ± 0.486pA, control, 10.64 ± 0.400pA, *p* = 0.009, unpaired T-test) (Figure 5A,B). The observed reduction in mEPSC amplitude aligns with the decrease in GluA1 surface density at the postsynaptic site that we observed by immunofluorescence (Figure 4C,D). Moreover, we detected a concordant decrease in the interevent cumulative percentage (Figure 5C; *D=* 0.595, *p <* 0.0001) and the frequency of interevent intervals in Ykt6 knockdown (0.357 ± 0.037 Hz) compared to the control (0.750 ± 0.010 Hz) (Figure 5D, *p*= 0.003, unpaired T-test with Welch’s correction). Importantly, the decrease in mEPSCs is not due to neuronal viability issues, as ATP levels—a reliable indicator of cell viability—remained unaffected by Ykt6 knockdown (Figure 5F).

**Figure 5.**
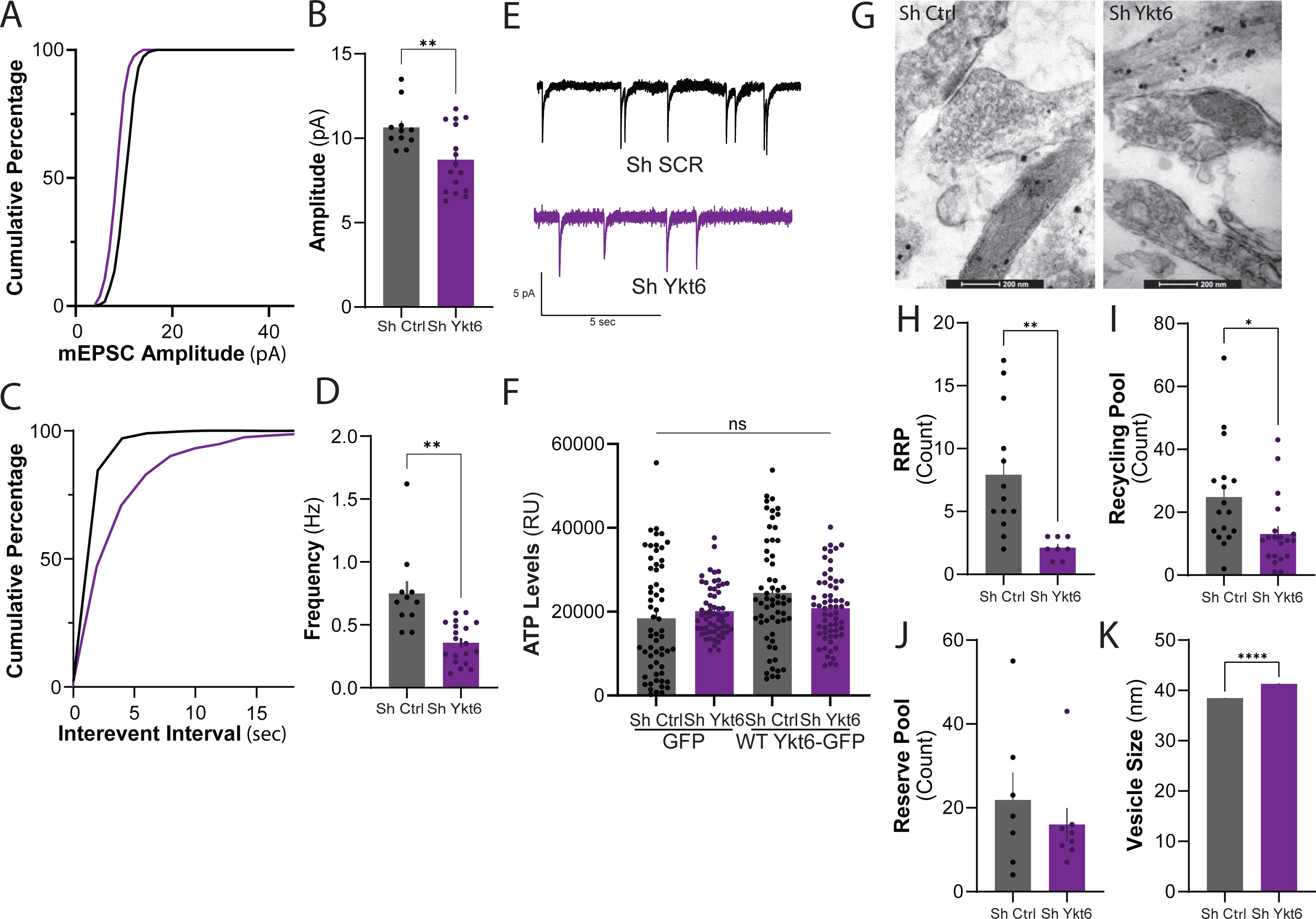
Ykt6 modulates both pre- and post-synaptic compartments of glutamatergic neurotransmission. Rat primary hippocampal pyramidal neurons were transduced with Sh Ctrl or Sh Ykt6 and patch-clamped in whole-cell configuration at DIV18-21 in the presence of tetrodotoxin (TTX), D-2-Amino-5-phosphonovalerate (D-APV), and picrotoxin (PTX) to isolate AMPA currents (A-D). **A)** Cumulative distribution of the amplitudes for all events per condition. **B)** Average amplitude per condition. **C)** Cumulative distribution of time between each event for every event per condition. **D)** Average frequency of events for each condition. (*p* < 0.0001, Kolmogorov-Smirnov test). **E)** Representative traces of cultures in (A); N=3, 4-6 cells per biological replicate. **F)** Quantitation of endogenous ATP levels at DIV21. N=3. **G)** Representative EM images of dendritic spines from the hippocampal cultures described above. **H)** Synaptic vesicle count for readily releasable pool (RRP). Each point is count for 1 image from (G). **I)** Synaptic vesicle count for recycling pool from (G). **J)** Synaptic vesicle count for reserve pool from (G). **K)** Average synaptic vesicle size from (G). All stats, Unpaired T-test with Welch’s correction, * p ≤ 0.05, ** p ≤ 0.01, **** p ≤ 0.0001.

To investigate whether the changes in synaptic transmission were due to changes in the number of vesicles associated with the synapse, we performed electron microscopy (EM) on rat primary pyramidal hippocampal neurons transduced with Sh Ctrl or Sh Ykt6. In line with previous studies (76, 77), we classified the vesicles that are touching synaptic contact points as the readily releasable pool (RRP), those proximal to the contact point but not touching membranes as recycling pool, and those distal from the contact point as reserve pool. Ykt6 knockdown exhibited spines with an immature filopodia-like morphology, whereas the control spines displayed a characteristic mature mushroom-like shape (Figure 5G). Furthermore, Ykt6 knockdown resulted in fewer synaptic vesicles in both the readily releasable pool (RRP) and the reserve pool, while the distal pool remained unaffected (Figures 5H–J). Additionally, we observed a slight increase in the average size of synaptic vesicles across all three pools in the Ykt6 knockdown condition (Figure 5K). Together, these data indicate that Ykt6 regulates the RRP, ultimately impacting mEPSC synaptic function in pyramidal hippocampal neurons.

### Ykt6 regulates dendritic arborization

The secretory pathway is not only important for neurotransmitter release and synapse morphology, but it also provides proteins essential for dendritic morphogenesis (97, 98). Dendritic arborization is crucial not only during neurodevelopment but also in adult neurons, where synaptic activity can influence the morphology of the dendritic arbor (39, 46, 78–83). To investigate whether Ykt6’s role in the secretory pathway affects dendritic morphology, hippocampal primary neurons transduced with either ShYkt6 or ShCtrl were transfected with mCherry to visualize the dendritic arbor and analyzed at DIV21 using Sholl analysis to assess branching complexity (84). Ykt6 knockdown reduced dendritic arbor branching in pyramidal hippocampal neurons compared to the control, particularly in regions distant from the soma (Figure 6A-C). Together, these data indicate that Ykt6 also plays critical role in the regulation of the dendritic arbor.

**Figure 6.**
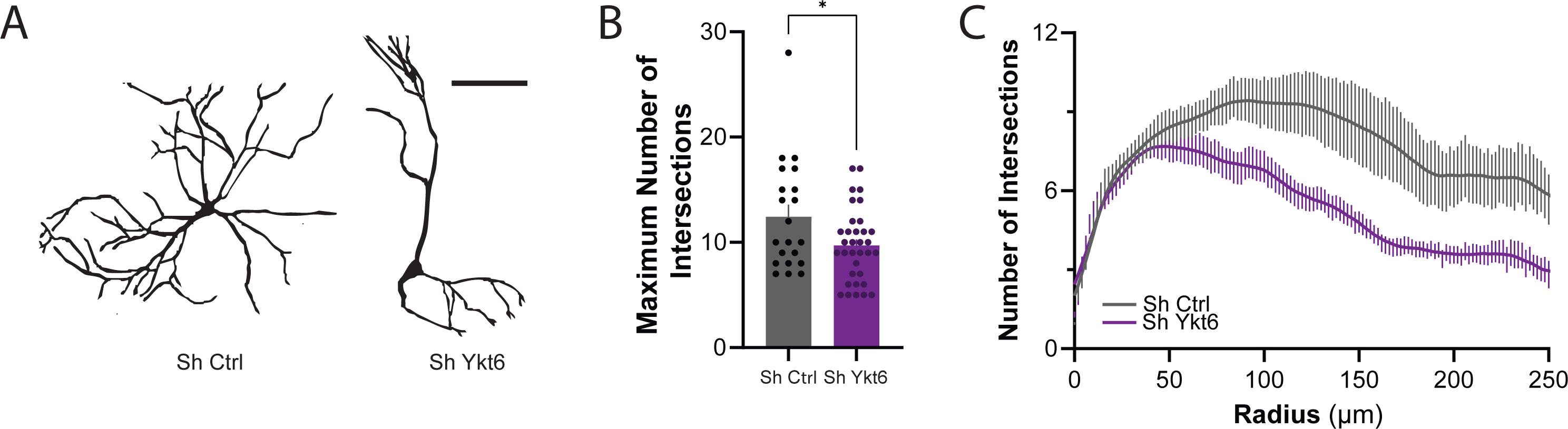
Dendritic arborization is dependent on Ykt6. Rat primary hippocampal neurons were transduced with either Sh Control or Sh Ykt6 RNA, transfected with PGK-mCherry construct on DIV19, immunoassayed for mCherry at DIV21 and analysed using Sholl analysis. **A)** Representative skeletonized images from pyramidal hippocampal neurons. **B)** Quantification of maximum number of intersections for hippocampal pyramidal neurons. **C)** Morphological analysis of hippocampal pyramidal neurons for the number of process intersections. Unpaired T-test with Welch’s correction. * p ≤ 0.05. N = 20-30, 10-15 cells per biological replicate. Scale bar: 100µm.

## Discussion

Ykt6’s role in vesicular trafficking has been well studied in yeast and mammalian systems, yet its physiological function in the brain remains largely unknown. Here, we show that Ykt6 is highly expressed in the mammalian hippocampus. In hippocampal pyramidal neurons, Ykt6 is associated with secretory organelles, ER and Golgi (105), and can be mobilized to postsynaptic terminals upon cLTP induction. Furthermore, we demonstrate that Ykt6 regulates both basal and cLTP-dependent surface GluA1 levels, modulating the availability of vesicular pools at the synapse and ultimately impacting mEPSCs, synapse structure, and dendritic morphology in pyramidal hippocampal neurons. Reductions in GluA1 expression may result from the presence of immature spines (85) which are also observed with the loss of Ykt6. Downregulation of Ykt6 not only disrupts GluA1 trafficking to the surface, reducing synaptic amplitude, but also contributes to longer inter-event intervals. These prolonged intervals have been linked to impairments in the readily releasable pool (86) and may also result from decreased spine density and the presence of immature spines (87, 88)—all of which we observe under Ykt6 deficiency.

Ykt6 is an essential R-SNARE protein with established roles in three key vesicular trafficking pathways: the secretory, exocytosis and autophagy pathways (1, 4–8, 10, 14, 15, 19, 89–95). Our findings suggest a novel role for Ykt6 in regulating AMPARs via the secretory pathway, distinct from its roles in autophagy or exocytosis. This hypothesis is supported by the following evidence. While surface expression of AMPARs can be modulated by autophagy in an activity-dependent manner (96–101), autophagy primarily facilitates the degradation of these receptors. In contrast, our data demonstrate that Ykt6 has a positive role in both basal and activity-dependent AMPAR trafficking, enhancing receptor surface expression rather than contributing to their degradation.

The R-SNARE VAMP2 (Synaptobrevin-2) plays a crucial role in forming well-established SNARE complexes, such as SNAP25–Syntaxin-1A/B–VAMP2 or VAMP2–Syntaxin-3–SNAP-47, which are essential for AMPAR exocytosis and endocytosis at postsynaptic terminals (102–108). Although Ykt6, another R-SNARE, could theoretically substitute for VAMP2 in these complexes, it remains unclear how Ykt6 might displace VAMP2 from its known binding partners. This suggests that Ykt6 is unlikely to regulate GluA1 surface expression via exocytosis.

Neurons have evolved specialized ER and Golgi outposts at distal locations from the soma to enable rapid and precise delivery of protein cargo to synapses (109, 110). Extensive networks of smooth ER tubules and sheets in dendrites and spines suggest local secretory trafficking, bypassing traditional somatic secretory pathways (28, 81, 111–116). ER exit sites (ERES) and COPII subunits have been observed at both proximal and distal dendritic locations in hippocampal neurons (112, 117, 118). Similarly, Golgi outposts in dendritic and axonal compartments support localized secretory functions essential for synaptic plasticity and neuronal responses. Local AMPAR synthesis in dendrites has been observed to rely on the ER-Golgi intermediate compartment (ERGIC) (119–121). Moreover, recent evidence suggests alternative sources of AMPAR synthesis coupled to synaptic plasticity-dependent signals (22, 122, 123). For example, in Drosophila, intra-Golgi transport is regulated by Ca²⁺-calmodulin-dependent kinase II, an enzyme activated during long-term potentiation (LTP) (124–127). Additionally, the dendritic ER plays a critical role in long-term depression (LTD), a form of synaptic plasticity. During LTD, ER Ca²⁺ release is enhanced, NMDAR trafficking is upregulated, and the ER network near spines becomes more The fact that Ykt6 is present at ER and Golgi outposts in secondary dendrites under basal conditions, and relocalizes to synaptic terminals upon cLTP induction, suggests a potential role of Ykt6 in distal secretory outposts. Understanding how Ykt6 regulates AMPAR secretory trafficking at distal secretory outposts will be crucial for elucidating its molecular mechanisms and functional significance, particularly in the context of neuronal plasticity and synaptic function.

Misfolding of α-synuclein leads to a group of neurodegenerative diseases collectively known as synucleinopathies, which includes Dementia with Lewy Bodies and Parkinson’s Disease Dementia (128–133). A block in ER-to-Golgi transport and a loss of Ykt6 function is a hallmark of α-synuclein pathology across various model systems (1, 4, 5, 17, 134, 135). Previous findings from our laboratory demonstrated that α-synuclein ER-to-Golgi trafficking deficits leads to a pathological increase in cytosolic Ca^2+^ levels and constitutive activation of the phosphatase CaN, leading to Ykt6 constitutive dephosphorylation and loss of function (1). Moreover, other groups have shown that α-synuclein disrupts AMPAR and NMDAR surface levels at synapses, impairing LTP and spatial learning (47, 49, 58, 136–138). Consistent with these findings, here we have shown that physiological Ykt6 loss of function in the hippocampus results in defects in both basal and cLTP-dependent GluA1 surface expression levels. Therefore, our data can provide a mechanistic explanation for how α-synuclein disrupts AMPAR trafficking and synaptic plasticity. Dysregulated AMPAR surface-level regulation has been implicated not only in Parkinson’s disease but also in other neurodegenerative disorders associated with cognitive and memory deficits, such as Alzheimer’s disease (122, 139–149), as well as a range of neurodevelopmental and psychiatric conditions. Therefore, our findings on the novel role of Ykt6 in AMPAR regulation within the hippocampus may open new therapeutic avenues for devastating neurological diseases.

### Experimental Procedures

#### Primary Hippocampal Cultures

Embryonic rat hippocampal neurons were isolated from euthanized pregnant Sprague–Dawley rats at embryonic day 18 using a modified protocol from Lesuisse and Martin. Protocol was approved by Northwestern University administrative panel on laboratory animal care. Embryos were harvested by Caesarean section and hippocampi were isolated and dissociated with 0.25% Trypsin without EDTA (Invitrogen, 15090-046) digestion for 15 min at 37°C and trituration with 1 ml plastic tip. Poly-D-Lysine (Sigma, P-1149)-coated 1.5 coverslips (Neuvitro, GG-12--1.5-oz) in 24-well plates were seeded with 5 x 10^5^ cells accordingly in neurobasal medium (Gibco, 21103-049) supplemented with 10% heat-inactivated FBS (Gibco), 0.5 mM glutamine (Gibco), penicillin (100 IU/mL), and streptomycin (100 μg/mL) (Gibco). Before seeding, cells were counted using the automated cell counter TC10 (Bio-Rad) and viability (90-95%) was checked with Trypan Blue Stain (0.4%, Gibco 15250-061). After 1 hour (h) incubation at 37°C, media was changed to neurobasal medium (Gibco, 21103-049) supplemented with B27 (Gibco, 17504-044) and 0.5 mM glutamine. One half (out of 500 µl volume for 24-well plates) of the media was changed on DIVs 5, 9, 12, 16, and 19.

#### Plasmids

Nontargeting ShRNA (Horizon Discovery Dharmacon, VSC11656) or ShRNA targeting Ykt6 (Horizon Discovery Dharmacon, V3SH7669) were expressed in mammalian cells using lentiviral induction at DIV 5.

Expression was verified visually by the expression of turboRFP.

Human eGFP-Ykt6 (a kind gift from Dr. Joseph Mazzulli, Northwestern University) and eGFP were expressed in mammalian cells using lentiviral induction at DIV 5 at half the MOI used with ShRNAs.

#### Doxycycline treatment of cultures

Primary hippocampal cultures that were infected with either nontargeting ShRNA or ShRNA targeting Ykt6 were treated with doxycycline. Targeted silencing of Ykt6 in primary hippocampal cultures was induced using doxycycline treatments at a concentration of 1nM at DIV8, 12, 15, and 19 before used for experiments. Viral MOI and doxycycline titrations were performed on new batches before applied experimentally and confirmation of knockdown and expression of turbo RFP was regulated.

#### Chemical LTP induction

The samples were washed once with warmed extracellular solution (ECS) containing (in mM): 150 NaCl, 2 CaCl_2_, 5 KCl, 10 HEPES, 30 Glucose, 0.001 TTX, 0.01 strychnine, and 0.03 picrotoxin at pH 7.4, then exposed to 300µM glycine in ECS for 3 minutes (external/internal staining) or 6 minutes (western blot) at room temperature, then washed with extracellular solution thrice, then incubated at 37°C for 20 minutes prior to further processing.

#### External/Internal Staining

External and internal glutamatergic receptors were stained according to the protocol by Chiu. et.al (72). Coverslips with primary hippocampal cultures were transferred to parafilm-covered dishes and incubated in primary antibody (GluA1: Invitrogen, MA5-27694; GluN1: Alomone Labs, AGC-001) at the concentration of 1:100 diluted in conditioned media for 15 minutes at room temperature, then washed three times with warm conditioned media. The coverslips were then washed once with phospho-buffered saline (PBS) supplemented with 1mM MgCl_2_ and 0.1mM CaCl_2_ (PBS+). The cells were then fixed by incubating with fresh 4% paraformaldehyde (PFA) and 4% sucrose in PBS for 8 minutes. The cells were then washed three times with PBS, then blocked with 10% normal goat serum (NGS) in PBS for minimum 30 minutes at room temperature. The cells were then incubated with 1:400 dilution of goat anti-mouse Alexa Fluor® 405 secondary antibody (Invitrogen, A-31553) or goat anti-rabbit Alexa Fluor® 405 secondary antibody (abcam, ab175652) in 3% NGS in PBS for 1 hour at room temperature. The cells were then washed 3 times with PBS.

For internal staining, the cells were then permeabilized with 0.25% Triton X-100 (Thermo Fisher Chemicals, A16046.AP) for 7 minutes at room temperature, then blocked with 10% NGS in PBS for minimum 30 minutes at room temperature. The intracellular receptors were then labelled by incubating the cells with the same primary antibodies at the same concentration, and anti-microtubule-associated protein 2 (MAP2) chicken antibody (abcam, ab318993) at 1:10000 dilution, but diluted in 3% NGS in PBS overnight in 4°C. The next morning, cells were washed 3 times with PBS, then labelled with goat anti-mouse Alexa Fluor® 647 secondary antibody (Invitrogen, A-21235) or goat anti-rabbit Alexa Fluor® 405 secondary antibody (Invitrogen, A-21245) at 1:1000 dilution and goat anti-chicken Alexa Fluor® 488 secondary antibody (Invitrogen, A-11039) at 1:400 dilution in 3% NGS in PBS for 1 hour at room temperature. The cells were then washed 3 times with PBS and once with de-ionized water, then mounted using Prolong Gold antifade mountant (Invitrogen, P36930). The cells were imaged using Nikon A1R GaAsP point-scanning laser confocal microscope using 63x oil-immersion objective (NAL=L1.4) with z-series of 10-20 images, taken at 0.3 μm intervals, with 2048x#x00D7;2048 pixel resolution. Only cells positive for MAP2 were acquired. The external to internal expression ratio was calculated for the basal condition with GFP and Sh Ctrl, or Sh Ctrl only. This value was then normalized to 1, and all other ratios normalized to the basal condition.

#### Ykt6 Staining

Primary hippocampal cultures as described above at DIV21 were washed once with PBS+, then fixed with fresh 4% PFA and 4% in sucrose in PBS for 20 minutes at room temperature. They were then permeabilized with 0.3% Triton-X 100 in PBS for 30 minutes at room temperature. The cells were blocked with 0.3% Triton-X 100/2% bovine serum albumin (BSA)/5% NGS in PBS for 30 minutes, then incubated for 48 hours at with mouse v-SNARE Ykt6p antibody (Santa Cruz, sc-365732), rabbit anti-Ykt6 primary antibody (abcam, ab236583), purified mouse anti-GM130 primary antibody (BD Biosciences, 610822) at 1:200 dilution, or rabbit anti-PDI primary antibody (Cell Signaling Technology, 3501), all at 1:100 dilution in the blocking buffer. Anti-microtubule-associated protein 2 (MAP2) chicken antibody (abcam, ab318993) at 1:10000 dilution was used to identify neurons. For PSD95, mouse anti-PSD95 primary antibody (Invitrogen, MA1-045) was used at 1:400 dilution.

After primary incubation, cells were washed three times with 0.05% Tween-20 in PBS for 15 minutes each, then with PBS twice for 15 minutes each, then incubated with then labelled with goat anti-mouse Alexa Fluor® 647 secondary antibody (Invitrogen, A-21235) or goat anti-rabbit Alexa Fluor® 405 secondary antibody (Invitrogen, A-21245) at 1:5000 dilution and goat anti-chicken Alexa Fluor® 488 secondary antibody (Invitrogen, A-11039) at 1:400 dilution in the blocking buffer for 2 hour at 4°C. The coverslips were then washed three times with 0.05% Tween-20 in PBS for 15 minutes each, and then twice in PBS for 15 minutes each, and mounted with Prolong Gold Antifade mountant. The cells were imaged using Nikon A1R GaAsP point-scanning laser confocal microscope using 63x oil-immersion objective (NAL=L1.4) with z-series of 10-20 images, taken at 0.3 μm intervals, with 2048x#x00D7;2048 pixel resolution. Only cells positive for MAP2 were acquired.

#### Sholl Analysis

Scholl Analysis was performed in cultured rat primary hippocampal cultures. Neurons were plated and allowed to mature until DIV21 and transfected with PGK-mCherry using Lipofectamine 2000 according to manufacturer’s protocol at DIV 16. At DIV21 they were washed once with PBS+, fixed with fresh 4% PFA and 4% in sucrose in PBS. They were then permeabilized with 0.3% Triton-X 100 in PBS for 7 minutes at room temperature, blocked with 3% NGS in PBS, and then stained with anti-mCherry primary antibody (Abcam ab167453) at the concentration of 1:400 diluted in 3% NGS in PBS. overnight in 4°C.

The next day, the coverslips were incubated with Goat Anti-Rabbit Alexa Fluor® 594 secondary antibody (abcam, ab150080) at 1:500 concentration diluted in 3% NGS in PBS, washed 3 times with PBS, and then mounted using Prolong Gold Antifade mountant. Isolated neurons were imaged using Nikon A1R GaAsP point-scanning laser confocal microscope using 63x oil-immersion objective (NAL=L1.4) with z-series of 8–10 images, taken at 0.3 μm intervals, with 1024L×L1024 pixel resolution. The stacks were then flattened to 1 image in ImageJ and Scholl Analysis performed using the SNT plugin (ImageJ)(150) after manual tracing.

#### Ykt6/Organelle Analysis

To measure the relative intensities of Ykt6 and the respective organelles, ImageJ was used with the ColorProfiler plugin. For the soma, a line was drawn in between the beginnings of the basal dendrites of the hippocampal neurons. For the dendrites, the line was drawn across the width of the secondary dendrite, immediately after the bifurcation point.

#### PSD95 distance measurement

To measure the distance between post-synaptic density 95 (PSD95) puncta and its most proximate Ykt6 puncta, coverslips were incubated per protocol above using the mouse anti-PSD95 and the rabbit anti-Ykt6 antibodies at given concentrations. The coverslips were imaged on Nikon AXR point-scanning laser confocal microscope using 100x (NA=1.49), with 2048x#x00D7;2048 pixel resolution and Nyquist. The images were analysed using ImageJ (84) and Colocalization by Cross Correlation package by McCall (151), by first constructing a mask to extrapolate the secondary dendrite, and then calculating the mean of all numerical values of the shortest distance of all PSD95 puncta with Ykt6 puncta.

#### Synaptosomal Fractionation

Synaptosomal fractionation was performed following a protocol adapted from Bermejo et.al (152). Adult brains were extracted in accordance with Northwestern University administrative panel on laboratory animal care post-decapitation. The brain was chopped with a razor blade and washed with ECS once before the glycine treatment. The brain was then snap frozen with liquid nitrogen until further processing.

The fractionation was performed as follows: solutions shown in table below were chilled, and HALT protease/phosphatase inhibitor (Thermo Scientific, 78440) was added at 1:100 dilution to all solutions.

**Table 1.**
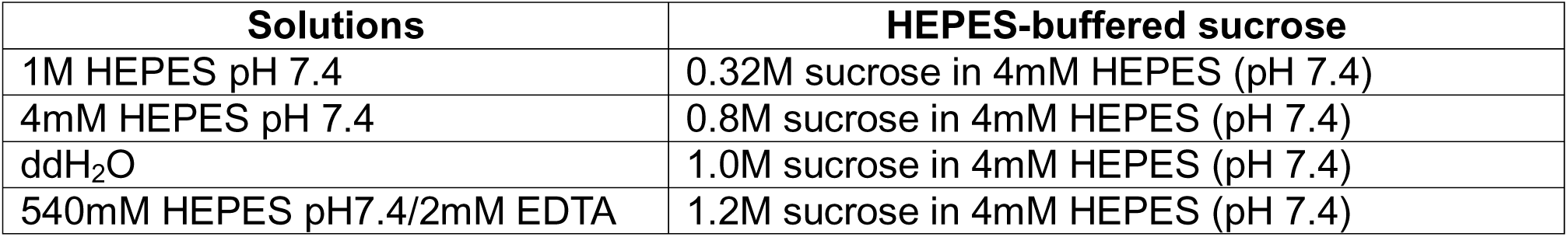
Reagents for synaptosomal fractionation.

All samples and solutions were kept on ice during the procedure. The tissue was placed in a dounce homogenizer along with 1mL of 0.32M HEPES-buffered sucrose solution and homogenized with the motor drive set at 900rpm (setting 7) over 30 second period with 12 strokes. The homogenizer was rinsed with deionized water and wiped dry with Kimwipes in between samples. 10µL of homogenate was preserved and diluted to 4 times the concentration using 0.32M HEPES-buffered sucrose solution, then stored at -80°C for subsequent protein quantification and western blot analysis of the total protein fraction.

The remaining homogenate was centrifuged in a fixed angle rotor at 900xg for 10 minutes at 4°C. The supernatant (S1) was transferred to a clean Eppendorf tube, while the nuclear fraction (P1) was resuspended in 500μL of 0.32M HEPES-buffered sucrose and stored at -80 °C and used for western blot analysis of the nuclear fraction.

S1 was then centrifuged at 10,000xg for 15 minutes at 4°C. The supernatant (S2) was removed and stored at - 80 °C. The pellet (P2) was resuspended in 1mL of 0.32M HEPES-buffered sucrose solution and centrifuged at 10,000xg for 15 minutes at 4°C. The supernatant (S2’) was removed and stored at -80 °C for subsequent protein quantification and western blot analysis of the cytosolic/light membrane fraction.

The pellet was then lysed by resuspending it in 1mL of ddH_2_O, transferred to a glass-Teflon tissue homogenizer, and rapidly homogenized by hand with 3 strokes. The sample was quickly adjusted to 4mM HEPES with 4μL of 1M HEPES solution, then returned to an Eppendorf tube, and the samples were then rotated at 4°C for 30 minutes to complete the lysing.

The sample was then centrifuged at 21,700xg for 24 minutes at 4°C. The supernatant containing the crude vesicular fraction (S3) was stored in -80°C, and the pellet was resuspended in 0.32M HEPES-buffered sucrose solution. The discontinuous sucrose gradient was prepared in 4mL open top thinwall ultra-clear tube (Beckman Coulter, 344062) by layering 1mL each of 1.2M HEPES-buffered sucrose solution, followed by 1.0M HEPES-buffered sucrose solution, then 0.8M HEPES-buffered sucrose solution using a P1000 pipette. Finally, the sample was layered on top, and the tubes were balanced using P200 pipette and adding 0.32M HEPES-buffered sucrose solution onto the top layer. The samples were then ultracentrifuged for 2 hours at 4°C in swinging buckets (Beckman Coulter, SW-60Ti) using Optima XE-90 ultracentrifuge (Beckman Coulter). Using an 18G needle and a 1mL syringe, the tubes were then punctured at the 1.0M/1.2M HEPES-buffered sucrose solution interphase and the band (synaptic plasma membrane layer, SPM) was withdrawn. The volume was noted, and the collected layer was placed in 3.5mL thickwall ultracentrifuge tubes (Beckman Coulter, 349622) and 2.5 times the volume of SPM of 4mM HEPES was added to adjust the sucrose concentration from 1.2M to 0.32M. The tubes were then balanced with 0.32M HEPES-buffered sucrose solution, and the samples were ultracentrifuged in the swinging bucket rotor at 200,000xg for 30 minutes at 4°C. The supernatant was discarded, and the pellet was resuspended in 100μL of 50mM HEPES/2mM EDTA solution, then stored in - 80 °C until used for western blotting.

#### Western Blot

Protein concentration was analyzed with the Pierce BCA Protein Assay kit (Thermo Scientific, 23225). After the addition of the appropriate amount of the 6X Laemmli Sample Buffer (Bio-rad sab03-02) with 5% ß-mercaptoethanol (Sigma Aldrich, M6250) protein samples were boiled and separated on precast 4-20% Criterion TGX Stain-free gels (Bio-Rad) and transferred to a nitrocellulose membrane (Amersham Protran 0.2 µm NC, #10600001). Membranes were blocked with 3% BSA in 1X Tris-buffered saline (TBS) (50mM Tris/Cl pH 7.4, 150mM NaCl) for 1 hour at room temperature. Membranes were subsequently immunoblotted overnight in primary antibody (anti-Ykt6: 1:100; anti-GAPDH: 1:500; anti-PSD95: 1:1000; anti-GluA1: 1:1000) at 4°C, shaking. The following day, membranes were washed three times with 1X TBST (TBS with 0.1% Tween) for 5 minutes and incubated in secondary IRDye antibody at 1:15000 dilution for 1 hour shaking at room temperature. Membranes were washed three times with 1X TBST for before imaging using Li-Cor Odyssey® CLx Imaging System. Images were processed and quantified using Image Studio Software (LI-COR Biosciences).

#### Electrophysiology

Mini EPSCs were recorded from infected primary hippocampal cultures in whole-cell voltage clamp. Recordings were performed between DIV 18-21. The external solution contained the following (in mM): 150 NaCl, 2.8 KCl, 2 CaCl_2_, 1 MgCl_2_, 10 glucose, and 10 m HEPES (pH adjusted to 7.3 with NaOH), with D-AP5 (50 µM), picrotoxin (50 µM), and tetrodotoxin (1µM) added. The internal solution used in recordings (in mM): 110 CsF, 30 CsCl, 10 Cs-HEPES, 5 EGTA, 4 m NaCl, and 0.5 CaCl_2_ (pH adjusted to 7.3 with CsOH). Cells were held at −70 mV in voltage clamp with an Axopatch 200B amplifier (Molecular Devices). Data were analyzed with Clampfit version 11.0.3 (Molecular Devices).

#### Electron Microscopy

Samples were prepared from an adapted protocol by Arai and Waguri (153). 10 minutes at room temperature and 50 minutes at 4°C. Cells were washed with 0.12 M phosphate buffer pH 7.4 and then treated with 1% osmium tetroxide and 1.5% potassium ferrocyanide (Sigma) in 0.12M phosphate buffer pH 7.4. Cells were dehydrated by an ascending series of alcohol (50, 70, 80, 90, and 100%) followed by treatment with epoxy resin for 24 hours. The grids were mounted on resin blocks and cured at 65°C for 3 days. The blocks were trimmed to contain CSMN before proceeding to ultra-thin sectioning. Resin blocks were sectioned on a Leica Ultracut UC6 ultramicrotome (Leica Inc., Nussloch, Germany). Sections (70nm) were collected on 200 mesh copper-palladium grids. Ultra-thin sections were counterstained on a drop of UranyLess solution (Electron Microscopy Sciences, Hatfield, PA) and 0.2% lead citrate. Grids were examined on FEI Tecnai Spirit G2 TEM (FEI Company, Hillsboro, OR) and digital images were captured on an FEI Eagle camera. Post-image acquisition adjustment was performed by Adobe Photoshop 2025.

Circular structures that were between 35 to 45 nm were considered as synaptic vesicles (77) and were manually counted. Those that were touching the membrane were counted as readily releasable pool (RRP), those close to the membrane but not touching were counted as recycling pool, and those distal to the membrane and the synaptic contact points were considered to be the reserve pool. Vesicular size was manually quantified using ImageJ.

### Statistical Analysis

Graphpad Prism 10 (http://graphpad.com) was used to graph, organize, and perform all statistical analysis. Statistical analysis was determined using the following methods: in case of more than two conditions, Bartlett’s Test was performed to test for homogeneity of variances. If the variances were heterogeneous, one-way analysis of variance (ANOVA) with Welch’s correction alongside with Dunnett’s T3 multiple comparisons test was performed; otherwise, standard ANOVA with multiple comparison was applied. In cases where two conditions were compared, F-test was performed to compare variances. If the variances were homogeneous, standard T-test was applied; otherwise, T-test with Welch’s correction was used to examine the significance of the results.

For the cumulative probability distribution, the event values were binned to construct the cumulative probability, and Kormagorov-Smirnov test was applied to examine whether the distribution differed significantly from each other.

## Data availability statement

All data are contained within the manuscript.

## Acknowledgements

We thank Geoff Swanson for kindly allowing us to use his electrophysiology rig, Farida Korobova from the Nikon Microscopy Center for her assistance with electron microscopy, and Anis Contractor and Adrian Contreras for their critical reviews.

## Funding and additional information

This study was funded by R01 NS117750 from the National Institute of Neurodegenerative Diseases and Stroke. The content is solely the responsibility of the authors and does not necessarily represent the official views of the National Institutes of Health.

## Conflict of Interest

The authors declare that they have no conflicts of interest with the contents of this article.

## Notes

### Competing Interest Statement

The authors have declared no competing interest.

### Summary of Updates

Introduction streamlined to focus more on glutamatergic transmission; discussion significantly changed to focus more on the role of Ykt6 in glutamatergic mechanisms; figure 2 revised to include all figures that were in the supplementary figure, lower part of figure 2 moved to figure 3.

